# Domain-specific folding of the tandem β-propeller protein Coronin 7 (Coro7) by CCT/TRiC

**DOI:** 10.1101/2025.03.11.642617

**Authors:** DeHaven J. McCrary, Teri Naismith, Silvia Jansen

## Abstract

The Chaperonin containing tailless complex polypeptide 1 (CCT) or TCP-1 ring complex (TRiC) plays a central role in maintaining cellular homeostasis by supporting protein folding and damping protein aggregation. Besides the abundant cytoskeletal proteins, actin and tubulin, CCT/TRiC is emerging as an obligate chaperone for WD40 proteins, which are comprised of one or multiple β-propeller domains. To date, only WD40 proteins consisting of a single β-propeller domain have been described as CCT/TRiC substrates. Using a combination of biotin proximity ligation, mass spec analysis and co-immunoprecipitation, we here identify the tandem β-propeller protein, Coronin 7 (Coro7), as a novel CCT/TRiC interactor. Transient knockdown of CCT/TRiC further severely diminished expression of Coro7, suggesting that Coro7 is a bona fide CCT/TRiC substrate. Interestingly, co-immunoprecipitation of truncated Coro7 proteins demonstrated that CCT/TRiC only interacts with the first β-propeller domain of Coro7. In line with this, fusion of a miniTurboID tag to the N- or C-terminus of Coro7 showed significant enrichment of all CCT/TRiC subunits for the first, but not the second β-propeller domain. Similarly, co-immunoprecipitation with individual Coro7 β-propeller domains generated by introduction of a protease cleavage site in full length Coro7, confirmed that CCT/TRiC only binds to the first β-propeller domain. Altogether, our study shows that CCT/TRiC can also function as a chaperone for multi-β-propeller domain proteins, likely by initiating the folding of the first β-propeller domain, which can then help template autonomous folding of consecutive β-propeller domains.

## INTRODUCTION

Correct protein folding is critical to maintain tissue homeostasis (1, 2). It ensures sufficient levels of proteins that are essential to sustain cellular function as well as prevents the excessive accumulation of misfolded and aggregation-prone proteins that could lead to the development of neuropathies and myopathies (3–6). The process of protein folding is initiated as soon as proteins emerge as simple strings of amino acids from ribosomes bound to the endoplasmic reticulum. However, whereas many smaller proteins will spontaneously assume their correct conformation, larger proteins with more complex folds will require the help of professional molecular folding machines consisting of a combination of chaperones, co-chaperones and chaperonins (7, 8). The latter comprise a family of proteins that require ATP hydrolysis to fold proteins inside of an enclosed chamber. In case of the group II TCP-1 ring complex (TRiC) (also called Chaperonin containing tailless complex polypeptide 1 or CCT), eight paralogous subunits (CCT1-CCT8) assemble into a 1 MDa hexadecameric double-ring structure (9–12). Each subunit consists of an apical domain that participates in substrate recognition, an equatorial domain that mediates ATP-hydrolysis and an intermediate domain that translates equatorial ATP hydrolysis into conformational changes of the apical domain (13, 14). As such, ATP hydrolysis following substrate binding results in repositioning of a flexible loop in the apical domain, thereby closing off the TRiC chamber and releasing the substrate in the chamber for assisted folding (12, 15). After ATP hydrolysis, the chamber opens and if the protein has achieved its proper fold and lost its contacts within the chamber, it is released.

The diverse amino acid composition of the CCT/TRiC apical domains creates a plethora of distinct hydrophilic and hydrophobic patterns that can recognize many different substrates. The most well-known CCT/TRiC substrates are the cytoskeletal proteins, actin and tubulin (16–18). In addition, CCT/TRiC seems to have a propensity for folding of WD40 or β-propeller domain proteins, including the heterotrimeric G-protein β-subunit (19–21), mTORC subunits mLST8 and Raptor (22) and the anaphase promoting complex proteins Cdc20 and Cdh1 (23, 24). A recent cryo-EM study was able to capture the stepwise folding of G-protein β5 (GNB5) inside the folding chamber of CCT/TRiC and elucidated the TRiC surfaces that guide this process (21). However, the different amino acid composition of CCT/TRiC β-propeller substrates calls into question whether they share a common CCT/TRiC recognition motif and whether the folding mechanism established for GNB5 is generally applicable to other CCT/TRiC β-propeller substrates. In addition, WD40 proteins in higher vertebrates can contain up to 4 β-propeller domains, whereas the folding chamber of CCT/TRiC has been estimated to only hold proteins of up to ∼70 kDa (25, 26) and thus can barely accommodate two β-propeller domains (∼ 40-50 kDa each). Given that no multi-β-propeller protein has been identified as a CCT/TRiC substrate to date, this raises the question whether mammalian CCT/TRiC is the designated chaperonin that mediates the folding of these complex proteins.

The extensive β-propeller family of the Coronins plays a prominent role in the remodeling of actin networks and hence is essential for many biological processes ranging from cell migration and intracellular transport to T-cell immune response and wound healing (27–29). In contrast to yeast, mammals have six different Coronin genes, which show ubiquitous or tissue-specific expression. In addition, Coronins underwent a gene duplication event, which led to the appearance of the class of the tandem Coronins as early as *Dictyostelium Discoideum* (*30*). As the name suggests, these Coronins are comprised of two consecutive Coronin β-propeller domains (**Fig. 1A**). They further lack the coiled-coil region present in the conventional Coronins and thus have been postulated to function as monomers rather than oligomers. Whereas the tandem Coronins from *Dictyostelium*, *C. Elegans* and *D. Melanogaster*, called CorB, and Pod-1 respectively, colocalize with F-actin, and are involved in actin-driven processes such as neurite outgrowth, polarity, endocytosis, and phagocytosis (31–35), it remains quite controversial whether these functions are conserved in mammalian Coro7. Instead, Coro7 seems to rather play a role as a scaffolding protein that supports the assembly of specific protein complexes. In addition, several genetic studies have identified Coro7 as a factor related to longevity, circadian rhythm and appetite (36–38), however it remains elusive how the scaffolding function of Coro7 contributes to regulating these processes.

**Figure 1:**
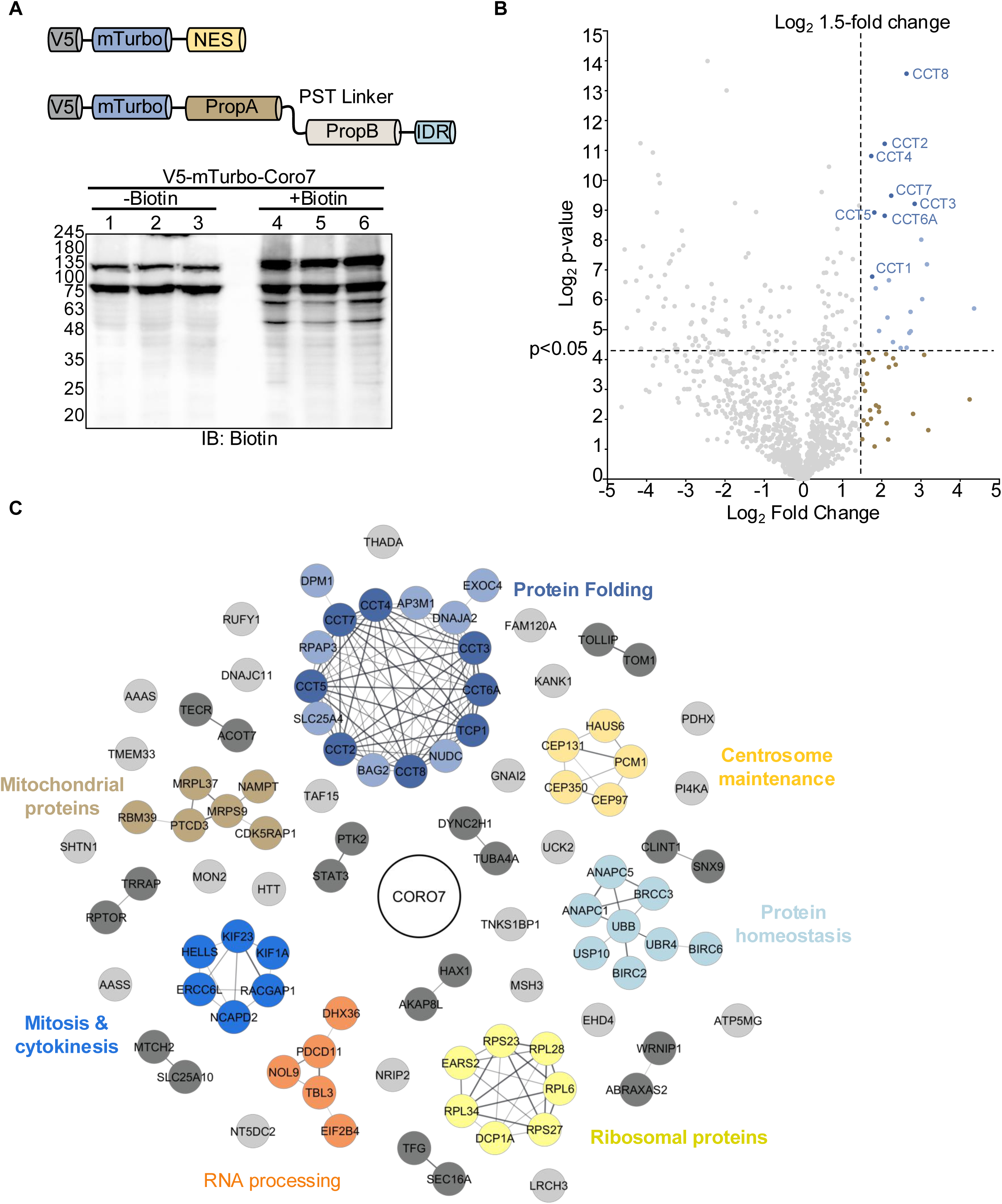
Biotin Proximity Ligation and mass spec analysis identify CCT/TRiC as a novel interactor of Coro7. **A)** Domain map of V5-mTurbo-NES cytoplasmic control and N-terminal V5-mTurbo tagged full-length Coro7 (V5-mTurbo-Coro7). Western blot using HRP-labelled streptavidin shows the increase in biotinylated proteins observed after addition of exogenous biotin to cells expressing V5-mTurbo-Coro7. **B)** Volcano plot showing all Coro7 interacting proteins identified by mass spec after proximity ligation using V5-mTurbo-Coro7. To identify significantly enriched proteins, peptide counts of each protein were Log_2_ transformed and compared to peptide abundance from the NES control group. Significant enrichment was gated at 1.5 Log_2_ fold change and a Log_2_ p-value of 4.322 (p ≥ 0.05). Data was analyzed from three independent experiments. Statistical significance was determined using an unpaired two-tailed Student’s t-test. **C)** MCL clustering analysis of proteins with >Log_2_ fold change of 1.0 over the NES control.

To begin to understand the cellular role of Coro7 and to expand upon the handful of Coro7 interactors that have been identified to date, we used biotin proximity ligation in combination with mass spec analysis to establish a Coro7 interactome. MiniTurboBioID analysis elucidated strong interaction between Coro7 and all eight subunits of the CCT/TRiC chaperonin complex, raising the question whether Coro7 supports the assembly of this complex. To investigate this, interaction between Coro7 and CCT/TRiC was first confirmed by immunoprecipitation of multiple CCT/TRiC subunits with both endogenous and recombinantly expressed Coro7. Interestingly, knockdown of CCT/TRiC significantly decreased the expression of Coro7, indicating that Coro7 does not function as a CCT/TRiC scaffold, but instead is a bona fide substrate of CCT/TRiC. As the first tandem β-propeller substrate of CCT/TRiC, we next sought to get a better understanding of how Coro7 is folded by CCT/TRiC. Surprisingly, analysis of truncated Coro7 proteins elucidated that CCT/TRiC interacts mainly with the first β-propeller domain of Coro7. This was further confirmed by comparison of the N-terminal and C-terminal miniTurboID interactome of Coro7, which only showed significant enrichment of CCT/TRiC when the miniTurboID tag was placed at the N-terminus of Coro7. In addition, full length Coro7 with a prescission protease cleavage site introduced between the β-propeller domains, only showed co-immunoprecipitation of CCT/TRiC with the first β-propeller in the presence of the protease. Altogether, our study establishes Coro7 as a novel CCT/TRiC substrate with a unique folding trajectory in which the first β-propeller domain is recognized and folded by CCT/TRiC, whereas the second β-propeller can fold independently of CCT/TRiC.

## RESULTS

### Identification of novel Coro7 interacting proteins by miniTurboBioID

To identify protein complexes that interact with Coro7, we built a Coro7 interactome using biotin proximity ligation in combination with mass spec analysis. We opted for proximity ligation based on the lack of discernable accumulation of Coro7 to specific organelles or vesicles (**Fig. S1**), which strongly suggests that most of its interactions are very dynamic and transient. To circumvent this problem, we used miniTurboID, which is a 3^rd^ generation biotin ligase that can rapidly and covalently attach a biotin molecule to proteins that are in close proximity of the protein of interest under cellular conditions (39). Potential interactors can then be specifically isolated using Streptavidin affinity purification and identified by a combination of mass spectrometry and proteomic analysis. To perform this experiment in a Coro7-free background, we first generated HEK293T Coro7 CRISPR/Cas9 knockout cells (HEK293T Coro7^-/-^) (See Materials and Methods). In addition, we developed an N-terminal miniTurboID fusion of human Coro7 (V5-mTurbo-Coro7) as well as a construct consisting of miniTurboID fused to tandem nuclear export signal motifs (V5-mTurbo-NES) to control for promiscuous interactions with highly abundant cytosolic proteins (**Fig. 1A**). Before performing the actual experiment, we optimized transient expression of these fusion proteins in HEK293T Coro7^-/-^ cells to obtain equivalent expression of V5-mTurbo-NES and V5-mTurbo-Coro7 as well as determined the optimal incubation time with biotin to obtain good labeling (**Fig. 1A and S2**). Using this optimized protocol, we generated three independent samples for each fusion protein and analyzed the biotinylated proteins by quantitative label-free mass spectrometry (**Fig. S2**). Potential Coro7 interacting proteins were identified by calculating the enrichment of individual proteins, which was obtained by comparing their relative abundance in the V5-mTurbo-Coro7 samples to the V5-mTurbo-NES control samples (40)

As expected, we observed considerable self-labeling of V5-mTurbo-Coro7. In line with the controversial role of Coro7 as an actin-regulatory protein, GO-analysis did not show enrichment of actin or actin-binding proteins. Instead, we identified a small cluster of proteins that are linked to the cell cycle and centrosome maintenance (**Fig. 1C**). Common to these potential interactors and the processes they are involved in is that they are all placed near the centrosome. Whereas this is in line with previous studies that reported perinuclear localization of endogenous Coro7 (41, 42), we mostly observed diffuse cytoplasmic staining of endogenous and recombinant Coro7 and only occasionally detected cells with perinuclear enrichment of Coro7 (**Fig. S1**). Together with their lower enrichment score (less than 2-fold), we decided to not follow up on these targets any further. Instead, we focused on clusters of proteins that all showed at least a 4-fold enrichment (Log_2_ fold change >2, p<0.05). Interestingly, this was observed for all eight subunits of the CCT/TRiC complex (**Fig. 1B-C**), which is the obligate chaperonin complex that folds actin and tubulin as well as several well-known β-propeller proteins. This raised the question whether Coro7 is a novel CCT/TRiC substrate or whether Coro7 perhaps functions as a scaffold that supports the assembly and function of CCT/TRiC, thereby indirectly regulating the cytoskeleton through folding of actin and tubulin by CCT/TRiC.

### Coro7 is a novel CCT/TRiC substrate

To further characterize the relation between Coro7 and CCT/TRiC, we first confirmed that endogenous and recombinant Coro7 interact with CCT/TRiC. To test binding under endogenous conditions, Coro7 was immunodepleted from HEK293T cell lysates and analyzed for bound CCT/TRiC by western blot. As shown in **Fig. 2A**, only beads coated with an antibody directed against Coro7 displayed a signal for the CCT5 or TCP1α (CCT1) subunit of CCT/TRiC, whereas beads coated with rabbit immunoglubulins (RIgG) did not. Similarly, transiently expressed EGFP or EGFP-Coro7 was pulled down from HEK293T cell lysates using EGFP-nanobody coated beads. When the resins were probed for the CCT2 and TCP1α subunits of CCT/TRiC by western blot, we again only observed a CCT/TRiC signal for beads carrying EGFP-Coro7, but not for beads decorated with EGFP (**Fig. 2B**). Given that the Coro7 and CCT2 antibodies were both raised in rabbit and that the size of the individual CCT subunits overlaps with that of the antibody heavy chain, we were unable to also probe for CCT2 in our endogenous Coro7 co-immunoprecipitation. To overcome this hurdle, we examined direct interaction of Coro7 and CCT/TRiC by sucrose gradient fractionation. For this purpose, HEK293T cell lysates were separated by centrifugation on a 5-40% sucrose gradient and collected as fractions spanning 3% sucrose increments. As expected, the majority of the CCT/TRiC complex, detected by using CCT2 as a proxy, was found in the higher density fractions (**Fig. 2C**). In addition, we observed a smaller amount of CCT2 signal in the lower density fractions, which is in line with the fact that CCT2 can also exist as a monomer (43, 44). When analyzing Coro7, we found that most of the protein was retained in the lower density fractions, however a faint, but clear band could also be seen in the higher density fractions (**Fig. 2C**). The distinct separation between these Coro7 populations and the lack of a Coro7 signal in the highest density fractions further indicated that the second Coro7 band does not correspond to aggregated Coro7. Together with the presence of a CCT2 signal in the same fraction, this instead strongly supports the idea that the higher density Coro7 band is part of a Coro7-CCT/TRiC supercomplex. Altogether, we conclude from these experiments that endogenous Coro7, like recombinant EGFP-Coro7, not only interacts with CCT1, CCT2 and CCT5, but can bind to the CCT/TRiC complex as a whole.

**Figure 2:**
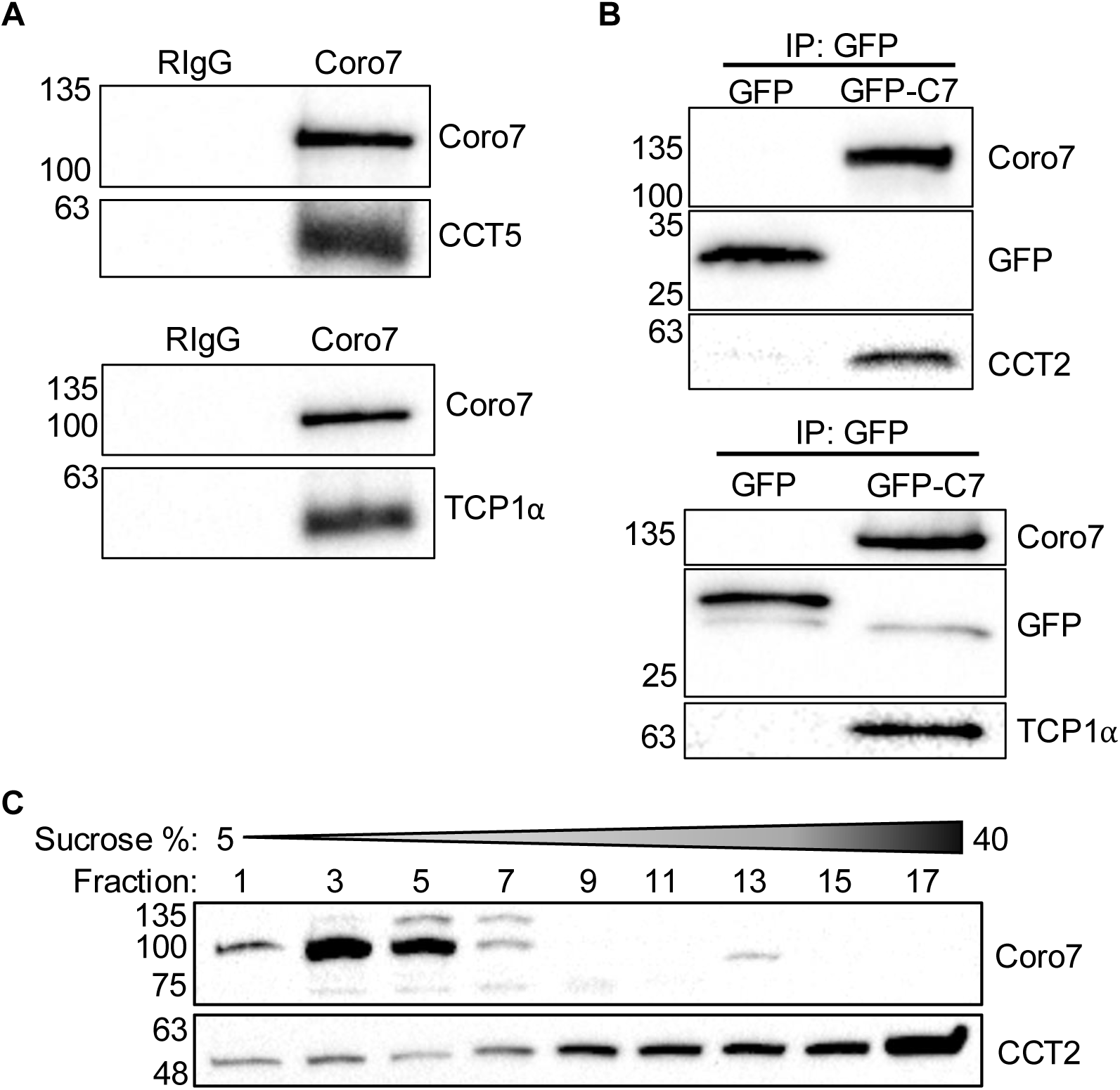
Endogenous and recombinant Coro7 interact with CCT/TRiC in HEK293T cells. **A)** Interaction between Coro7 and CCT/TRiC was confirmed by co-immunoprecipitation of CCT/TRiC subunits, CCT5 and TCP1α, with Coro7 or rabbit immunoglobulins (RIgG) as control. **B)** Co-immunoprecipitation of CCT/TRiC subunits, CCT2 and TCP1α with recombinant EGFP or EGFP-Coro7. **C)** Western blot showing Coro7 and CCT/TRiC after sucrose gradient fractionation (5-40%) of whole cell HEK293T lysates. Coro7 was present in two distinct populations in fraction 3-5 and fraction 13, whereas CCT/TRiC was mostly detected in higher sucrose fractions. Each fraction corresponds to a 3% increase in sucrose concentration.

Having established that CCT/TRiC and Coro7 are binding partners, we next investigated the biological role of this interaction. To examine whether Coro7 is a novel substrate of CCT/TRiC and requires the chaperone for its folding, we examined the effect of CCT/TRiC depletion on the protein expression level of Coro7. As it was previously shown that depletion of one subunit is sufficient to disrupt the formation of the CCT/TRiC complex, we worked with a mix of siRNAs directed against the CCT2 subunit, whereas control cells were transfected with a non-targeting scrambled siRNA (SCR). Western blot analysis showed a complete loss of CCT2 in CCT2 siRNA treated cells compared to control cells (**Fig. 3A-B**). In addition, these cells also showed a lower CCT5 signal, suggesting that the CCT/TRiC complex is indeed disrupted (**Fig. 3A**). Knockdown of CCT/TRiC further decreased Coro7 protein levels by about 50% (p < 0.0001) (**Fig. 3B**), suggesting that Coro7 is indeed a novel CCT/TRiC substrate.

**Figure 3:**
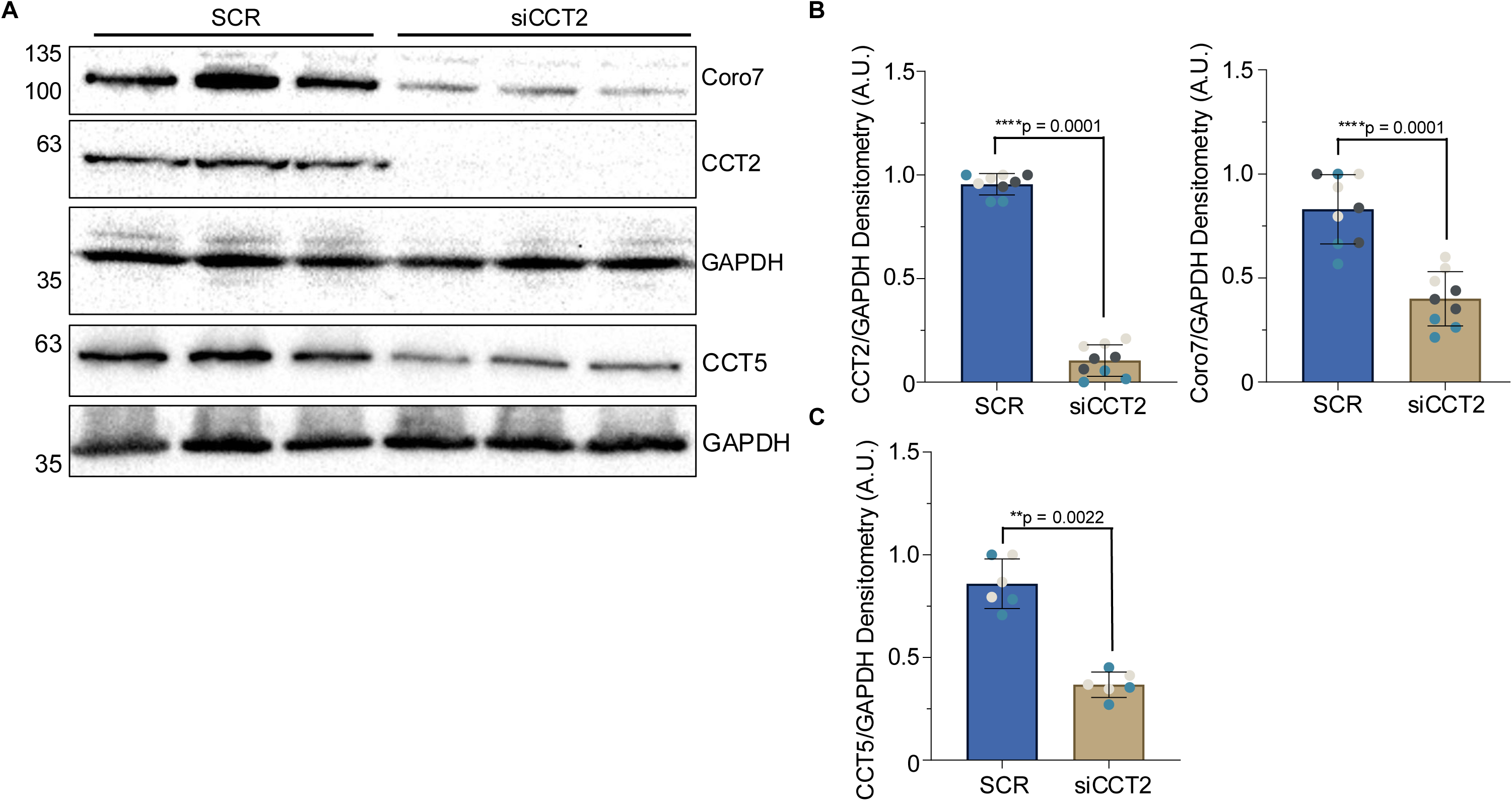
Interaction of Coro7 with CCT/TRiC is necessary for its biosynthesis. **A)** Western blot showing transient knockdown of CCT2 in MDA-MB-231. **B)** Densitometry analysis of CCT2 and Coro7 expression from scrambled and siCCT2 treated MDA-MB-231 cells. Data was analyzed from three independent experiments performed in triplicate for each condition for each repeat (N=3, n=3). Technical replicates are indicated by similar colored data points. **C)** Densitometry analysis of CCT5 expression from scrambled and siCCT2 treated MDA-MB-231 cells (N = 2, n = 3). Statistical significance was determined using a a non-parametric Mann-Whitney test. Exact p-values are shown in each graph.

### Only the first β-propeller of Coro7 interacts with CCT/TRiC

In contrast to other CCT/TRiC β-propeller substrates identified to date, Coro7 consists of two consecutive β-propeller domains and hence is also larger than the 70 kDa that can be accommodated by the CCT/TRiC folding chamber. This raised the question whether CCT/TRiC perhaps folds each β-propeller sequentially or whether CCT/TRiC has a different mechanism to induce the folding of either β-propeller domain. To address this, we generated a series of truncated Coro7 proteins. Due to the lack of a structural model of Coro7, truncation constructs were developed based on multiple sequence alignment of Coro7 with the canonical Coronins (Coro1-6) and by the predicted Coro7 structure in Alphafold. This enabled us to not only delineate both β-propeller domains, but also demonstrated that the proline-serine rich linker between the β-propellers as well as the C-terminal region of Coro7 are largely unstructured. Interestingly, the latter sequence is not found in any of the other Coronins except for yeast Crn1, where it was recently shown to be an intrinsically disordered region (IDR) that regulates the function of Crn1 (45). In line with this, analysis of the Coro7 sequence by Metapredict (46) demonstrated that the C-terminal region does indeed classify as an IDR (**Fig. 4B**) and from hereon we will refer to it as such.

**Figure 4:**
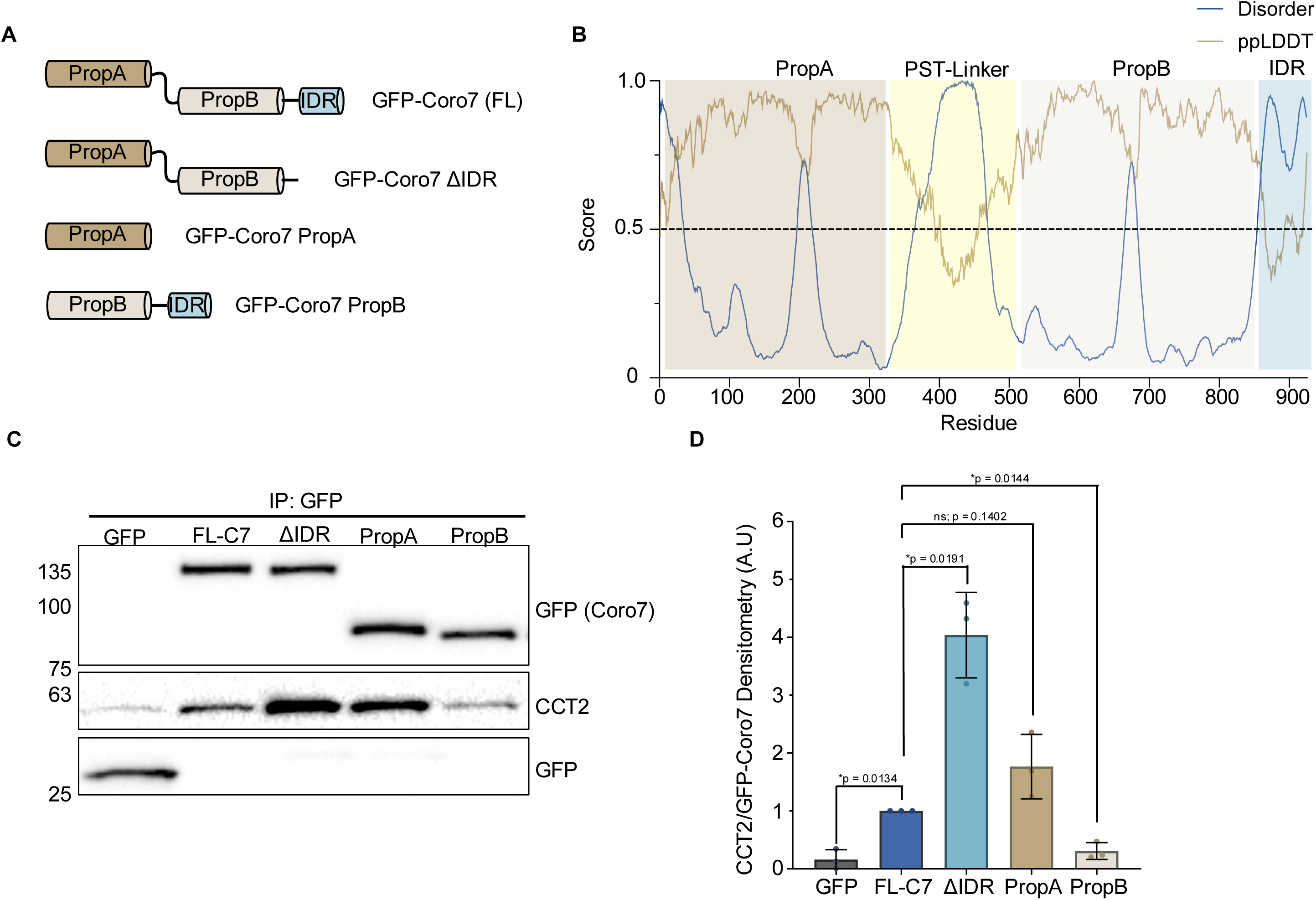
CCT/TRiC interacts with the first propeller of Coro7 in the absence of other domains. **A)** Domain maps of truncated Coro7 constructs. Amino acid residues are indicated for each construct. **B)** Metapredict v3.0 prediction of intrinsically disordered regions of full-length Coro7. Regions with disorder scores above 0.5 are predicted to be disordered with 50% confidence. The pLDDT score reflects the confidence of AlphaFold2 in the local structure prediction. **C)** Interaction between CCT/TRiC and different Coro7 domains was examined by co-immunoprecipitation of CCT2 with recombinant EGFP or EGFP-tagged Coro7 truncation constructs. **D)** Densitometry analysis of CCT2 precipitated by the indicated truncated Coro7 protein. Data was analyzed from three independent experiments. Statistical significance was determined using a Welch’s t-test comparing each truncated Coro7 with full length Coro7. Exact p-values are reported with exception of p>0.05, which is not considered significant (ns).

Based on this information, we generated EGFP-tagged Coro7 truncation constructs consisting of either the first (PropA) β-propeller, the second (PropB) β-propeller together with the C-terminal IDR or both propellers without the C-terminal IDR (ΔIDR) (**Fig. 4A**). After transient expression, truncated Coro7 proteins were immunoprecipitated from HEK239T cells using an EFGP-nanobody and analyzed by western blot for bound CCT/TRiC. Surprisingly, we found that PropA and Coro7-ΔIDR pulled down CCT/TRiC, whereas PropB barely showed a CCT2 signal compared to the EGFP only control (**Fig. 4B**). On top of that, densitometric analysis further showed that Coro7-ΔIDR precipitated significantly more CCT/TRiC than full length Coro7 (**Fig. 4C**). Altogether, these results unexpectedly elucidated that CCT/TRiC is only required for folding of the first β-propeller domain and that there might be an additional role for the C-terminal IDR to efficiently release Coro7 (PropA) from the CCT/TRiC folding chamber.

To examine whether this interaction pattern is similar for full length Coro7, we leveraged the short working distance (within 10 nm of the bait protein) of the miniTurboID tag and repeated biotin proximity ligation with an N-terminal as well as a C-terminal miniTurboID fusion of Coro7, referred to as V5-mTurbo-Coro7 and V5-Coro7-mTurbo, respectively (**Fig. 5A**). Next, we analyzed the enrichment of the CCT/TRiC subunits in either sample compared to the V5-mTurbo-NES control. CCT1-CCT8 were found in both the N- and C-terminal biotin ligation reaction, however whereas each subunit showed increased binding for V5-mTurbo-Coro7 compared to the control sample, only CCT1 and CCT3 were slightly enriched for interaction with V5-Coro7-mTurbo, while CCT2, CCT5 and CCT6 binding was downregulated (**Fig. 5B**, data points shown in red). This difference became even more clear when directly examining the enrichment of each CCT subunit in the N-terminal versus the C-terminal Coro7 interactome, which showed lower interaction of each subunit, except CCT3, for the C-terminal than the N-terminal miniTurboID Coro7 fusion (**Fig. 5C**). Proximity ligation analysis thus supports the observation that CCT/TRiC mainly interacts with the first β-propeller.

**Figure 5:**
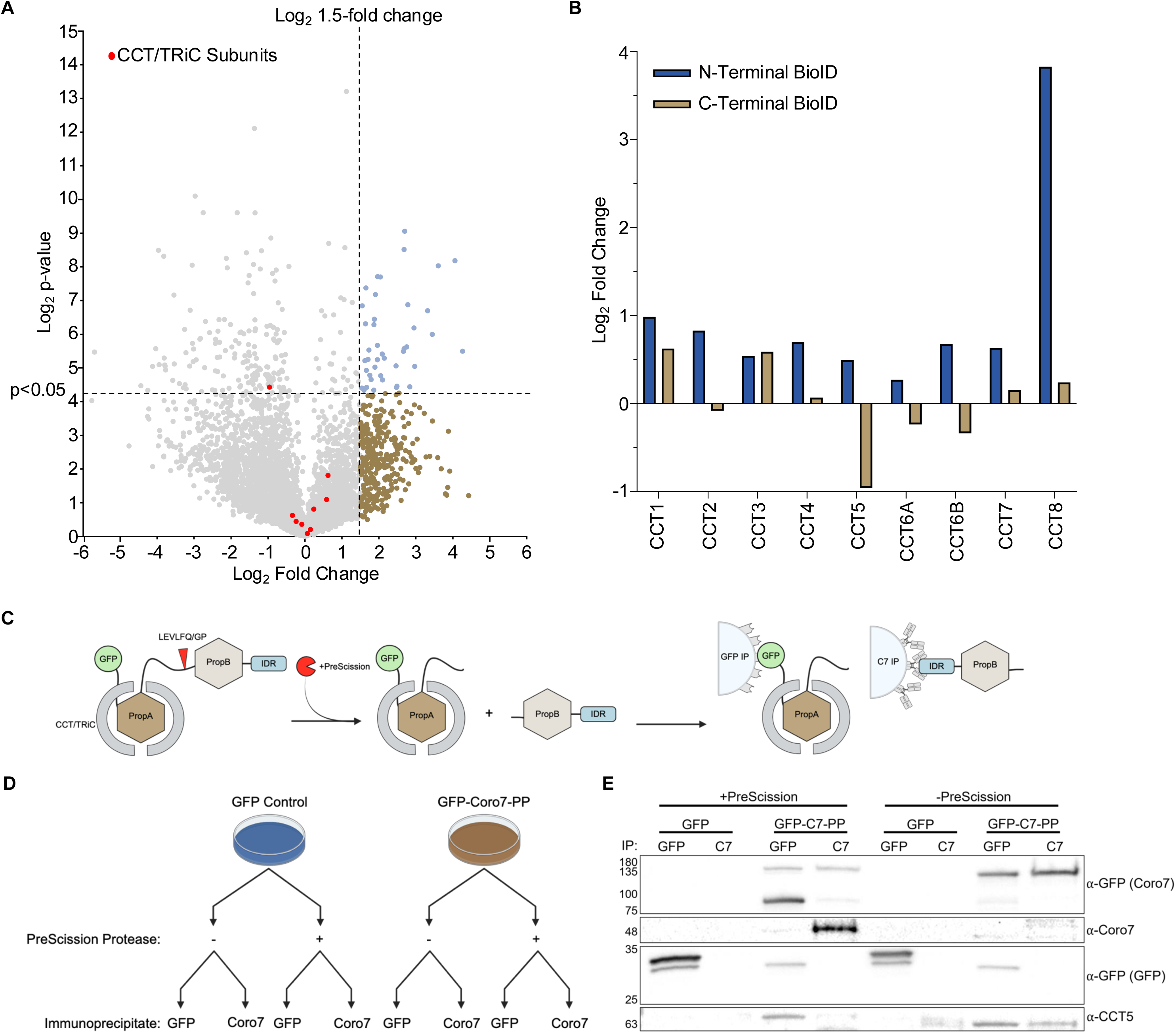
Domain-specificity of CCT/TRiC is conserved in full length Coro7. **A)** Volcano plot showing all Coro7 interacting proteins identified by mass spec after proximity ligation using C-terminal v5-mTurbo-Coro7. Significantly enriched proteins were identified as in Figure 1B. CCT/TRiC subunits (CCT1-8) are indicated in red. Data was analyzed from three independent experiments. Statistical significance was determined using an unpaired two-tailed Student’s t-test.. **B)** Direct comparison of enrichment (shown as Log_2_ fold change) of individual CCT/TRiC subunits from biotin proximity ligation assays using N-terminal and C-terminal fusion of V5-mTurboID and Coro7. **C)** Schematic overview of EGFP-Coro7-PreScission-Protease cleavage and subsequent co-immunoprecipitation of CCT/TRiC using either EGFP-nanobody or Coro7 antibody coated resin. **D)** Overview of sample acquisition workflow. One 10cm dish of HEK293T Coro7 KO cells transfected with either pEGFP-C1 or pEGFP-Coro7-PreScission Protease is split into 4 samples and subjected to the outlined treatment. **E)** Western blot showing co-immunoprecipitation of CCT/TRiC subunit CCT5 with EGFP and Coro7 in the presence and absence of prescission protease.

In addition, we generated full length EGFP-Coro7 with a prescission protease cleavage site introduced in the flexible linker between the β-propeller domains to more directly assess CCT/TRiC binding to either propeller. For this purpose, co-immunoprecipitation was performed using beads coated with either EGFP-nanobody or an antibody raised against the C-terminal 50 amino acids of Coro7 to specifically pull down the first or second β-propeller domain (**Fig. 5D**). In the absence of prescission protease, only intact Coro7 was detected and a signal for CCT5 was observed after co-immunoprecipitation with either pulldown strategy. By contrast, in the presence of the protease, no signal was observed for intact Coro7 and bands corresponding to either EGFP-PropA or PropB appeared (**Fig. 5E**). In line with our previous results, CCT5 only co-immunoprecipitated with EGFP-PropA, but not PropB (**Fig. 5E**). Altogether, this demonstrates that only PropA of Coro7 interacts with CCT/TRiC, suggesting that only the first β-propeller domain of Coro7 requires assisted folding whereas the second β-propeller domain folds autonomously or with the help of another interacting protein.

## DISCUSSION

The mammalian chaperonin, TRiC, has been estimated to fold >10% of the cytosolic proteome. This is mainly accounted for by its indispensable role in the folding of the abundant cytoskeletal proteins, actin and tubulin. In addition, CCT/TRiC is required for the folding of a growing list of proteins with a WD40 or β-propeller domain. Using a combination of biotin proximity ligation. mass spec analysis, co-immunoprecipitation and siRNA knockdown, we show here that the tandem β-propeller protein, Coro7, is a novel substrate of CCT/TRiC (**Fig 1-3**), elucidating for the first time that CCT/TRiC also participates in the folding of WD40 proteins that consist of multiple β-propeller domains.

β-propeller domains are generally comprised of seven WD40 repeats that fold into seven β-sheets, which are arranged around a central axis like the blades of a propeller (47). β-propeller domains further display another unique feature where the N-terminus folds back to complete the final β-sheet of the last blade that closes the propeller, referred to as the molecular clasp or velcro (48, 49). It has been postulated that this specific structural organization is the reason why β-propeller proteins require CCT/TRiC to complete folding. In line with this, a recent cryo-EM study was able to capture snapshots of the folding trajectory of GNB5 by CCT/TRiC (21). This work elucidated that CCT/TRiC actively directs the folding of the β-propeller structure by establishing specific intramolecular interactions that drive folding from the middle blade (blade 4) fanning out to the outer blades (blades 3 and 5, then blades 2, 6 and 7) to finally induce the first blade that will close the propeller. This is different in proteins that consist of multiple β-propeller domains, where the N-terminus folds back to complete the last blade in the neighboring propeller rather than close the first propeller and additional stabilization is obtained by interaction with other domains or by atypical blades (50–54).

Coro7 is predicted to be a tandem β-propeller protein and thus has been speculated to assume the same closed protein fold as other known tandem β-propeller domain proteins (55). Our study shows here that only the first β-propeller domain of Coro7 binds to CCT/TRiC (**Fig. 4C-D** and **Fig. 5**), raising the question how PropB is folded to the point that the N-terminus of Coro7 can insert itself to complete the tandem β-propeller. Given our observation that the IDR of Coro7 promotes the release of Coro7 from CCT/TRiC (**Fig4. C-D**), it is tempting to speculate that PropB is stabilized by another interaction partner that is recruited by the IDR until the tandem propeller structure can be closed by the Coro7 N-terminus. In respect to this, it should be noted that the C-terminal Coro7 interactome did not contain any known CCT/TRiC co-chaperones, such as Hsp70 (56). This is quite surprising, as Hsp70 preferentially binds to linear sequences that will assemble into β-sheets (56). However, Hsp70 contact with emerging β-sheets is generally short-lived and thus it is possible that Hsp70 molecules have dissociated by the time the C-terminal miniTurbo-ID tag becomes folded and active.

Instead of co-chaperones, we did find many targets that are WD40 proteins, suggesting that PropB might be stabilized by a β-propeller domain from another protein. This is particularly interesting as the canonical Coronins (Coro1-6) consist of only one β-propeller domain, but have an additional coiled-coil domain that enables them to form trimers (57–59). What is more, Coronins need to form oligomers in order to bind to actin filaments and exert their cellular functions (60). As Coro7 already has two β-propeller domains and lacks a coiled-coil domain, it is possible that it fulfills this requirement by interacting with other WD40 proteins. As we detected WD40 proteins involved in different processes, this might explain the diffuse cytoplasmic localization of recombinantly expressed Coro7 in contrast to the reported Golgi localization of endogenous Coro7 (41, 42). Going forward, it will be interesting to test whether WD40 binding partners can induce localization of Coro7 to the Golgi or even increase colocalization with F-actin.

Our observation that only one β-propeller of Coro7 interacts with CCT/TRiC is further in line with previous work showing that CCT/TRiC handles substrates that do not fit its folding chamber by only enclosing the domain of the protein that requires assisted folding (61). It does raise the question how CCT/TRiC recognizes β-propeller domains that need support during folding in contrast to β-propeller domains that can fold autonomously or with the help of other proteins. Using G-protein β as a model protein, Kubota and colleagues (62) showed that hydrophobic residues in β-sheets are critical for recognition by CCT/TRiC. Ribosome profiling of nascent WD40 domains expanded upon this notion by determining that CCT/TRiC rather recognizes folding intermediates like β-sheets than linear sequences (56). Currently, we have no structural information on Coro7 beyond an AlphaFold prediction, which counterintuitively folds PropA and PropB as two independent β-propeller domains instead of a closed tandem β-propeller structure. Until we have a better insight into the 3D organization of the Coro7 β-propeller domains, it will thus remain elusive which specific elements in PropA versus PropoB are recognized by CCT/TRiC.

Finally, our initial intent was to identify new interaction partners of Coro7 to gain insight into its enigmatic correlation with longevity, appetite and regulation of circadian rhythm. Instead, we identified Coro7 to be a novel substrate of CCT/TRiC, whose expression and activity have been reported to decline with aging (7), leading to disruption of proteostasis and build-up of protein aggregates. The decrease in Coro7 expression upon transient depletion of CCT/TRiC (**Fig. 3**) without concomitant decline in tubulin or actin levels (not shown), suggests that Coro7 and its potential WD40 interaction partners might be among the first CCT/TRiC substrates that get affected by even slightly reduced CCT/TRiC activity. As such, it will be interesting to determine if there is a correlation between CCT/TRiC and Coro7 expression in tissues of different ages and relate this to changes in metabolism, the onset of senescence and altered expression circadian clock genes in cells derived from these tissues.

## ACKNOWLEDGEMENTS

This project was supported by NIGMS R01 GM138448 and a Global Incubator Seed Grant from the WashU Office of the Provost to SJ. Mass Spectrometry analyses were performed by the Mass Spectrometry Technology Access Center at the McDonnell Genome Institute (MTAC@MGI) at Washington University School of Medicine, supported by the Diabetes Research Center/NIH grant P30 DK020579, Institute of Clinical and Translational Sciences/NCATS CTSA award UL1 TR002345, and Siteman Cancer Center/NCI CCSG grant P30 CA091842.

## AUTHOR CONTRIBUTIONS

DJM designed and performed experiments; TN performed experiments; SJ conceptualized amd designed research, wrote paper.

## MATERIALS AND METHODS

### Reagents

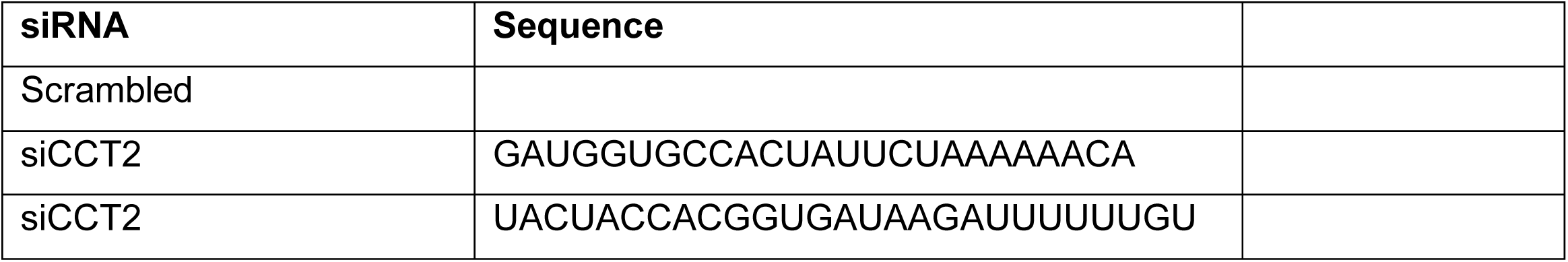
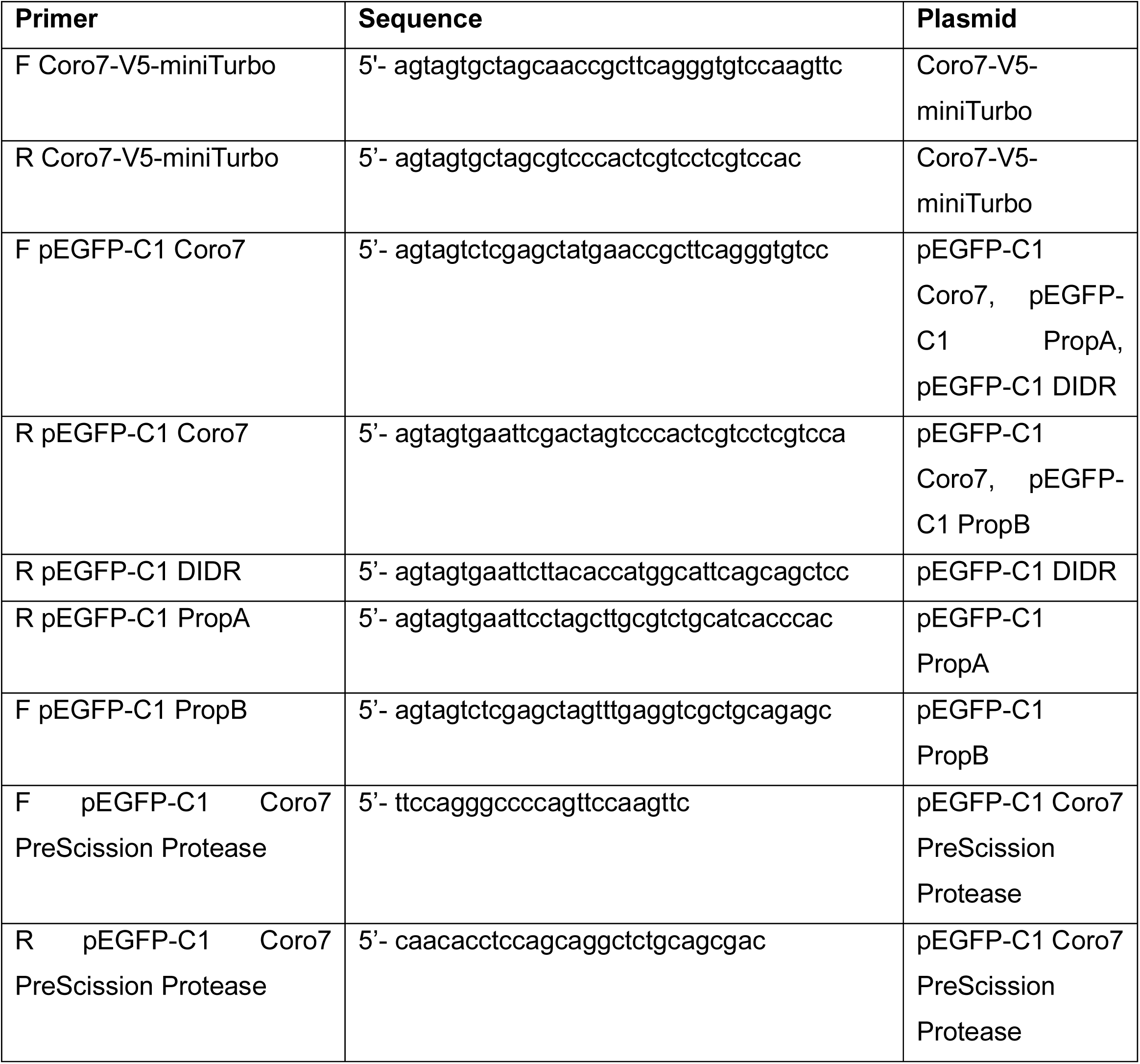

| Antibody | Company & Catalog | Dilution | Species |
| --- | --- | --- | --- |
| Coro7 | Novus #NBP3-18451 | 1:1000 (WB); 1:200 (IF) | Rabbit |
| Coro7 | Rybakin *et al.*, 2004 | N/A (IF) | Mouse |
| GFP | Santa Cruz #sc-9996 | 1:1000 (WB) | Mouse |
| TCP1a | Santa Cruz #sc-53454 | 1:1000 (WB) | Rat |
| CCT2 | CST #3561S | 1:1000 (WB) | Rabbit |
| CCT5 | Novus #NBP2-43680 | 1:1000 (WB) | Mouse |
| V5-tag | CST #13202S | 1:1000 (WB) | Rabbit |
| a/b-Tubulin | CST #21485 | 1:1000 (WB) | Rabbit |
| Streptavidin-HRP | Leinco #S207 | 1:1000 (WB) | N/A |
| GAPDH | Santa Cruz #sc-365062 | 1:1000 (WB) | Mouse |
| Rabbit IgG Control | Invitrogen #02-6102 | N/A | Rabbit |
| Anti-mouse IgG-HRP | CST #7076S | 1:10000 (WB) | Horse |
| Anti-rabbit IgG-HRP | CST #7074S | 1:10000 (WB) | Horse |
| Anti-rat IgG-HRP | CST #7077S | 1:10000 (WB) | Goat |
| Goat-anti-rabbit, Oregon Green 488 | ThermoFisher #O11038 | 1:500 (IF) | Goat |
| Donkey-anti-mouse AlexaFluor 488 | ThermoFisher # A21202 | 1:500 (IF) | Donkey |

#### Plasmid Cloning

A complete ORF encoding for human Coro7 was obtained from the DNASU plasmid repository (pDNR227) and cloned into V5-miniTurbo-NES (gift from X). For V5-miniTurbo-Coro7, Coro7 was cut out of an existing construct with AgeI and XhoI and ligated into V5-miniTurbo-NES using KpnI and XhoI restriction sites, destroying the AgeI and Kpn2I sites after ligation. For Coro7-V5-miniTurbo, Coro7 was generated by PCR and cloned into the NheI restriction site. Insertion and correct orientation of Coro7 was determined by colonyscreen. EGFP-tagged Coro7 and Coro7 truncation constructs were generated by PCR and cloned into pEGFP-C1 (Clontech) using XhoI and EcoRI restriction sites. Insertion of a PreScission Protease, recognition sequence was generated using Q5 mutagenesis. All constructs were confirmed by sequencing. Primers used to generate the plasmids used in this study are shown in Reagents section above.

#### Mammalian Cell Culture

HEK293T WT and Coro7 KO, MDA-MB231 WT and Coro7 KO, and ARPE19 WT cells were cultured at 37°C and 5% CO_2_ in DMEM, high glucose, GlutaMax (Gibco) supplemented with 10% fetal bovine serum (FBS) (Biowest #S1620) and 1% penicillin-streptomycin solution (Gibco).

#### Coro7 CRISPR/Cas9 KO HEK239T cells

100µM of tracrRNA and crRNA were mixed and heated at 95°C for 5 minutes to create a tracrRNA-cRNA duplex. This duplex was cooled to room temperature and combined with Cas9 in sterile PBS. The RNA and Cas9 were incubated together for 20 minutes at room temperature to form a ribonucleoprotein (RNP). To introduce the RNP to HEK293T cells were electroporated using a Lonza 4D Nucleofector. Briefly 2×10^5^ HEK293T WT cells were resuspended in 20uL of Nucleofector Solution SE. Next, 5µL of RNP and 1µL of Cas9 Electroporation Enhancer was then added to the resuspended cells. After mixing, the cells were then transferred to a Nucleocuvette strip and electroporated using the DG-130 program. After electroporation, cells were transferred to pre-warmed DMEM containing 10% FBS. To generate monoclonal populations, electroporated HEK293Ts were seeded as single cells in 96-well plates, allowed to form colonies and then harvested for Next Generation Sequencing.

#### Biotin Proximity ligation and mass spec analysis

1.5×10^6^ HEK293T Coro7 KO cells were seeded onto 10cm plates one day prior to transfection. HEK293T Coro7 KO cells were transfected with either 0.75µg of V5-mTurbo-NES (cytoplasmic control) or 3µg of V5-mTurbo-Coro7 plasmid using PEI per manufacturer’s instructions (Polysciences). 48 hours after transfection, cells were treated with 500µM exogenous biotin for 10 minutes at 37°C then lysed in 350µL of 20mM Tris-HCl pH 7.4, 150mM NaCl, 1% Triton X-100, 1X protease inhibitor cocktail (APeXBIO) and 1mM PMSF lysis buffer. Lysates were incubated on ice for 30 minutes with agitation every 10 minutes. Lysates were clarified by centrifugation at 21,000xg for 20 minutes and the supernatant was collected. 200µL of Dynabeads MyOne Strepatvidin T1 beads (ThermoFisher) were equilibrated by washing three times in 500µL of lysis buffer. Following equilibration, lysates were added to the equilibrated Dynabeads and incubated at room temperature for 2 hours under gentle agitation. The beads were then washed twice with RIPA-LS (10mM Tris-Cl pH 7.4, 1mM EDTA, 140mM NaCl, 0.1% SDS, 0.1% DOC, and 1% Triton X-100), once with 1M KCl, once with 1M Na_2_CO_3,_ and twice with 2M urea in 10mM Tris-Cl pH 7.4. The final wash was removed, and the beads were resuspended in 30µL of 3X Laemmli buffer. 5µL of sample was separated by SDS-PAGE and stained with Coomassie to check for protein pulldown. The remaining 25µL of sample was loaded onto a Mini-Protean TGX Stain-Free Gel, 4-20% (Bio-Rad) and run into the stacking gel. Following electrophoresis, the gel was briefly washed in water and samples were cut out of the gel in 1cm x 0.5cm pieces. Gel pieces were washed in water twice for 10 minutes each. Samples were processed for mass spectrometry by the Washington University in St. Louis School of Medicine Mass Spectrometry Technology Access Center (MTAC). Protein fold change and p-values were calculated as previously described by Aguilan *et al.,* 2020 (40). Briefly, raw spectrum counts for both the NES control and Coro7 experimental groups were subjected to a Log_2_ transformation to generate a normalized skew.

Following transformation, the Log_2_ spectrum counts were first normalized by the average of all spectrum counts in each control group and further normalized by the distribution width or slope of each control group. Next, missing values were imputed to account for missing data due to detection limitations. Finally, the fold change of Log_2_ spectrum counts in the Coro7 experimental group was calculated compared to the average of the NES control spectrum counts. p-values were determined using a two-tailed, paired t-test. Log_2_ fold changes were then plotted against the Log_2_ calculated p-values. For Markov Clustering algorithm (MCL) of enriched proteins (fold change >Log_2_ 1.0), protein-protein interactions were identified using STRING v12.0 (63) against the Homo sapiens protein database with a 0.400 confidence interaction score. Protein networks were then exported to Cytoscape v3.10.3 (64) where the network was clustered using MCL with a granularity parameter of 4.

#### Western Blotting

For Western blotting experiments, 1.5×10^6^ HEK293T WT or KO cells were seeded onto 10cm plates one day prior to transfection. Cells were transfected with either 2µg of pEGFP-C1 empty vector control or pEGFP-C1 Coro7 constructs (WT ΔCA, PropA, or PropB) using PEI (Polysciences) per manufacturer’s instructions. Transfections proceeded for two days prior to lysis. Cells were lysed in 20mM Tris-HCl pH 7.4, 150mM NaCl, 1% Triton X-100, 1X protease inhibitor cocktail (APeXBIO) and 1mM PMSF. Lysates were incubated on ice for 30 minutes with agitation every 10 minutes, then clarified by centrifugation at 21,000 x g for 20 minutes. The supernatant was collected and mixed with 3X Laemmli buffer and heated at 95°C for 5 minutes. Samples were separated by 12% SDS-PAGE and proteins were transferred onto Immobilon-P polyvinylidene difluoride (PVDF) membranes (Millipore Sigma) in transfer buffer containing 10% ethanol. Transfers proceeded for 1 hour at 100V. Membranes were allowed to dry and were blocked in 5% BSA in TBS plus 0.01% Tween-20. Blots were incubated with indicated primary antibodies at 1:1000 dilution overnight at 4°C with gentle agitation. Following primary incubation, blots were washed with TBS-T and incubated with horseradish peroxidase (HRP)-linked anti-mouse or anti-rabbit IgG secondary antibodies for 1 hour at room temperature with gentle agitation. Blots were washed with TBS-T and incubated in Clarity Western ECL substrate (Bio-Rad) for at least 1 minute before imaging. Blots were imaged using a BioRad ChemiDoc Imager.

#### Purification and conjugation of HIS-GFP Nanobody

This HIS-GFP nanobody was a gift from Dr. Brett Collins at the University of Queensland in Brisbane Australia. 50ng of HIS-GFP Nanobody cDNA was transformed into 50µL of BL21 competent cells and colonies were grown overnight on carbenicillin-containing agar plates. Starter cultures were inoculated using single colonies and grown in 100mL of LB at 37°C at 215 rpm. The following day, 1 liter TB cultures were inoculated and grown at 37°C at 215 rpm for until the OD600 was ≥ 2. Once desired cell density was reached, cultures were cooled to 18°C and induced with 500µM IPTG and protein production proceeded overnight. The following day, cultures were harvested by centrifugation at 3000xg for 10 minutes and lysed by microfluidization in 200mL of 20mM Tris-Cl pH7.4, 500mM NaCl, 10mM Imidazole, 4mM Benzamidine, 1mM PMSF, and 1X protease inhibitor cocktail (APeXBIO). Lysates were clarified by centrifugation at 12,000 x g for 30 minutes. Supernatants were then passed over 4mL bed volume of NiNTA equilibrated 3x in 20mM Tris-Cl pH 7.4, 500mM NaCl, and 10mM Imidazole by gravity flow twice. The NiNTA was washed with 20CV of 20mM Tris-Cl pH 7.4, 500mM NaCl, and 10mM Imidazole and bound protein was then eluted in 10 2mL fractions with 20mM Tris pH 7.4, 50mM NaCl, and 250mM Imidazole. Protein-containing fractions were determined by Coomassie stain, combined, and concentrated to a 10mL volume using an Amicon Ultra 3000Da MWCO spin column (Millipore). Concentrated protein fractions were then passed through a HiLoad 26/600 Superdex 75pg size exclusion column equilibrated in 200mM NaHCO_3_ and 500mM NaCl. Proteins were eluted using an isocratic elution over 1CV at a flow rate of 1.5mL/minute and a 0.2CV void volume. Peaks were checked by Coomassie stain and HIS-GFP Nanobody containing fractions were combined and concentrated to 2.5mL using an Amicon Ultra 3000Da MWCO spin column (Millipore). The HIS-GFP Nanobody was desalted into 3.5mL of PBS using a PD-10 desalting column (GE Healthcare). The HIS-GFP Nanobody was then diluted to 13mL and added to 1 gram of Pierce NHS-activated dry agarose (Thermo Scientific). NHS-nanobody conjugation proceeded overnight at 4°C with gentle agitation. The following day, the NHS beads were washed 3 times with PBS and the reaction was quenched using 1M Tris-Cl pH 7.4 for 15 minutes.

#### GFP Pulldowns

HEK293T Coro7 KO cells were transiently transfected with either 2µg of pEGFP-C1 (empty vector), or 4µg of pEGFP-C1-Coro7 WT, pEGFP-C1-Coro7-ΔCA, pEGFP-C1-Coro7 PropA, or pEGFP-C1-Coro7 PropB using a 1:3 DNA to PEI ratio (Polysciences). Following transfection, cells were lysed in 20mM Tris-HCl pH 7.4, 150mM NaCl, 1% Triton X-100, 1X protease inhibitor cocktail (APeXBIO) and 1mM PMSF. A 50µL bed volume of GFP-Nanobody beads were equilibrated in lysis buffer. Lysates were passed over the GFP-Nanobody beads once and beads were subsequently washed three times in 20mM Tris-HCl pH 7.4 and 150mM NaCl. After the final wash, the GFP-Nanobody beads were pelleted and resuspended in 75µL of 3X Laemmli sample buffer and heated to 95°C. The GFP-nanobody beads were then pelleted by centrifugation at 21,000xg for 1 minute and the supernatant was collected and used for Western blotting.

#### Endogenous Coro7 Co-IPs

1.5×10^6^ HEK293T cells were seeded onto 10cm dishes the day prior to Co-IP. Cells were lysed in 200µL of 20mM Tris-HCl pH 7.4, 150mM NaCl, 1% Triton X-100, 1X protease inhibitor cocktail (APeXBIO) and 1mM PMSF. Lysates were incubated on ice for 30 minutes with agitation every 10 minutes, then clarified by centrifugation at 21,000xg for 20 minutes. Following clarification, 2ug of either rabbit IgG or anti-Coro7 antibody were added to cell lysates and incubated at 4°C for 1 hour under gentle agitation. Protein-antibody complexes were isolated by the addition of a 25µL bed volume of magnetic protein G beads (NEB) for 1 hour under gentle agitation. The magnetic protein G beads were washed with 3x with 20mM Tris-HCl, pH 7.4 and 150mM NaCl and protein was eluted by the edition of 50µL of 3X Laemmli buffer and heating at 95°C for 5 minutes. The magnetic protein G beads were pelleted by centrifugation at 21,000xg for 1 minute and the supernatant was collected and used for Western blotting.

#### Sucrose Gradients

1.5×10^6^ HEK293T cells were seeded onto 10cm dishes the day prior to lysing. Cells were lysed in 350µL of 20mM Tris-HCl pH 7.4, 150mM NaCl, 1% Triton, 1X protease inhibitor cocktail (APeXBIO) and 1mM PMSF. Lysates were incubated on ice for 30 minutes with agitation every 10 minutes, then clarified by centrifugation at 21,000xg for 20 minutes. Sucrose gradients were formed by the mixing of 5% and 40% sucrose in 20mM Tris-Cl, pH 7.4 and 150mM NaCl. 11mL gradients were deposited using a Labconco Auto Densi-Flow Gradient Fractionator. Lysates were gently layered on top of cold 5-40% sucrose gradients and centrifuged at 26,300 RPM at 4°C for 16 hours. Following separation, 0.5mL fractions were taken and subsequently used for Western blotting.

#### siRNA Knockdown of CCT2

2.5×10^5^ MDA-MB-231 WT cells were seeded into a 6-well plate the day prior to siRNA transfection. MDA-MB-31 WT cells were transfected with either 20nM of scrambled siRNA or siRNA targeting CCT2 (see table for siRNA sequences) using Lipofectamine RNAiMAX Transfection Reagent (ThermoFisher) following manufacturer’s instructions. Briefly, 6 pmol of either scrambled or CCT2 siRNA was diluted in 150µL of OptiMEM. In a separate tube, 5µL of RNAiMAX was diluted in 150µL of OptiMEM. siRNA containing tubes were mixed with RNAiMAX containing tubes, and incubated at room temperature for 10 minutes. Following incubation, siRNA-RNAiMAX complexes were added dropwise to cells in antibiotic-free DMEM supplemented with 10% FBS and GlutaMAX. The following day, cell media was replaced with fresh complete media. siRNA knockdown proceeded for 72 hours and cells were then lysed, and lysates were used for subsequent Western blotting as previously described.

#### GFP-Coro7-PreScission Protease Cleavage and Co-IPs

1.5×106 HEK293T Coro7 KO cells were seeded onto 10cm dishes the day prior to transfection. Either an empty pEGFP-C1 vector or pEGFP-Coro7-PreScission-Protease (GFP-Coro7-PP) were transfected using PEI at a 1:3 DNA to PEI ratio (Polysciences). Following transfection, cells were lysed in 500μL 20mM Tris-HCl pH 7.4, 150mM NaCl, 1% Triton X-100, 1X protease inhibitor cocktail (APeXBIO) and 1mM PMSF. Lysates were incubated on ice for 30 minutes with agitation every 10 minutes. Lysates were clarified by centrifugation at 21,000xg for 20 minutes. GFP-Coro7-PP was then cleaved by the addition of 50µg of PreScission Protease for 1 hour at 4°C. Following cleavage, lysates were split in half with one half subjected to GFP pulldowns and the other half subjected to Coro7 pulldowns as described above. Protein was eluted by the addition of 3X Laemmli sample buffer and heated to 95°C and used for subsequent Western blotting

#### Lipofectamine 3000 Transfection

The day prior to transfection, 0.5×10^5^ MDA-MB-231 WT or Coro7 KO cells were seeded onto glass bottom D35 imaging dishes. Cells were transfected with Lipofectamine 3000 at a 1:4 ratio. Briefly, in one tube, 2µg of GFP-Coro7 and 0.5µg of BFP-LifeAct plasmid, and 5µL of P3000 reagent were diluted in 125µL of OptiMEM. In a separate tube, 10µL of Liopfectamine 3000 was diluted in 125µL of OptiMEM. The tubes were then combined with each other, incubated at room temperature for 15 minutes and added dropwise to the cells. The following day, the media was changed to fresh DMEM supplemented with 10% FBS.

#### Immunofluorescence Staining and Microscopy

0.5×10^5^ MDA-MB-231 WT or ARPE19 WT cells were seeded onto circular 12mm #1.5 glass coverslips the day before fixation. Cells were briefly washed with warm PBS and fixed for 10 minutes in pre-warmed 4% PFA in PBS at room temperature. Following fixation, cells were washed in ice-cold PBS. Cells were then permeabilized by the addition of 0.3% Triton X-100 in PBS for 10 minutes at room temperature, followed by a PBS rinse. Coverslips were then blocked in 3% BSA in PBS + 0.1% Tween (PBS-T) for 1 hour at room temperature. Primary antibodies were diluted in 3% BSA in PBS-T at indicated dilutions (see table) and incubated on coverslips for 1 hour at room temperature. Following primary antibody incubation, coverslips were washed 3 times in PBS-T for 5 minutes each. Secondary antibodies and Alexa Fluor 647-Phallodin were diluted in PBS-T at indicated dilutions (see table) and incubated on coverslips for 1 hour at room temperature in the dark. Coverslips were then washed 3 times in PBS-T for 5 minutes each. Finally, coverslips were mounted on to glass slides using ProLong Glass Antifade Mountant (Thermo Fisher) after briefly dipping in DIH_2_O to remove excess salts. Fixed slips and overexpressing live cells were imaged using a Nikon Ti2 inverted microscope with a ×100 Plan-Apo oil immersion objective and a Yokogawa CSU-W1 spinning disk confocal attached to a Hamamatsu ORCA-FLASH4.0 CMOS camera. Images were at 16-bit in a 2048×2044 resolution with Z-stacks of 0.2µm. Images were processed using Fiji to create sum-intensity projections.

---

**Figure S1:**
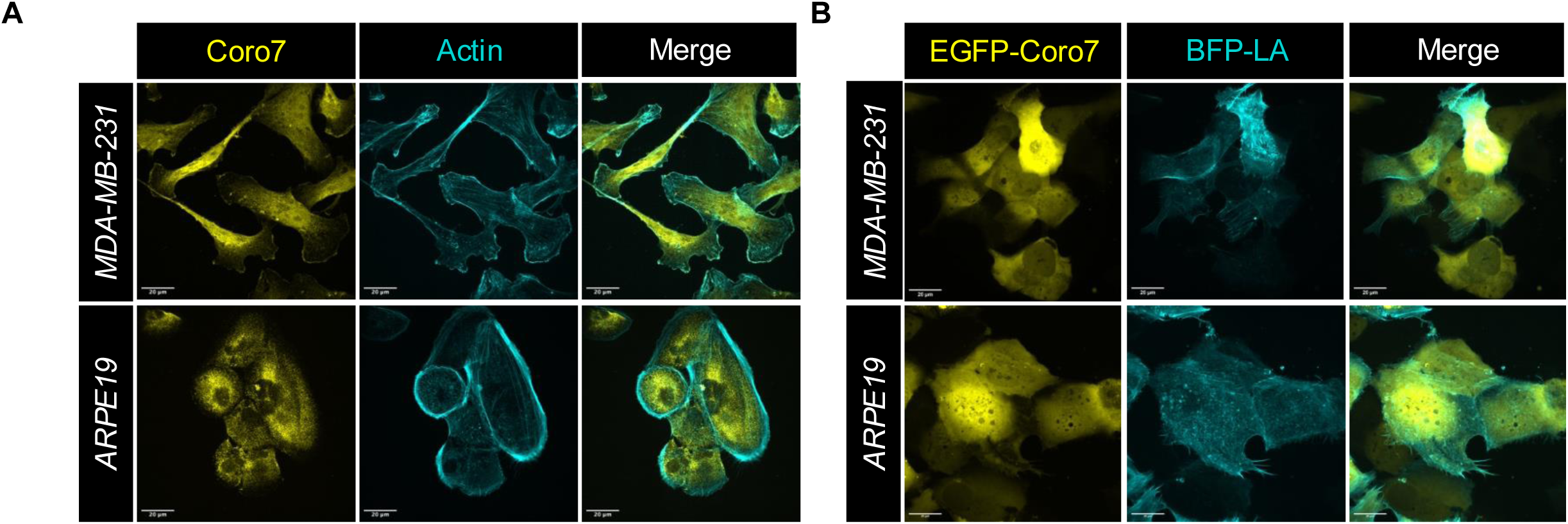
Coro7 mainly exists as a cytoplasmic protein. **A)** Confocal fluorescence images of MDA-MB-231 and ARPE19 cells immunostained for endogenous Coro7 and phalloidin (F-actin). **B)** Confocal live-cell images of MDA-MB231 and ARPE19 cells transiently expressing EGFP-Coro7 and BFP-LifeAct (BFP-LA). Scale = 20µm

**Figure S2:**
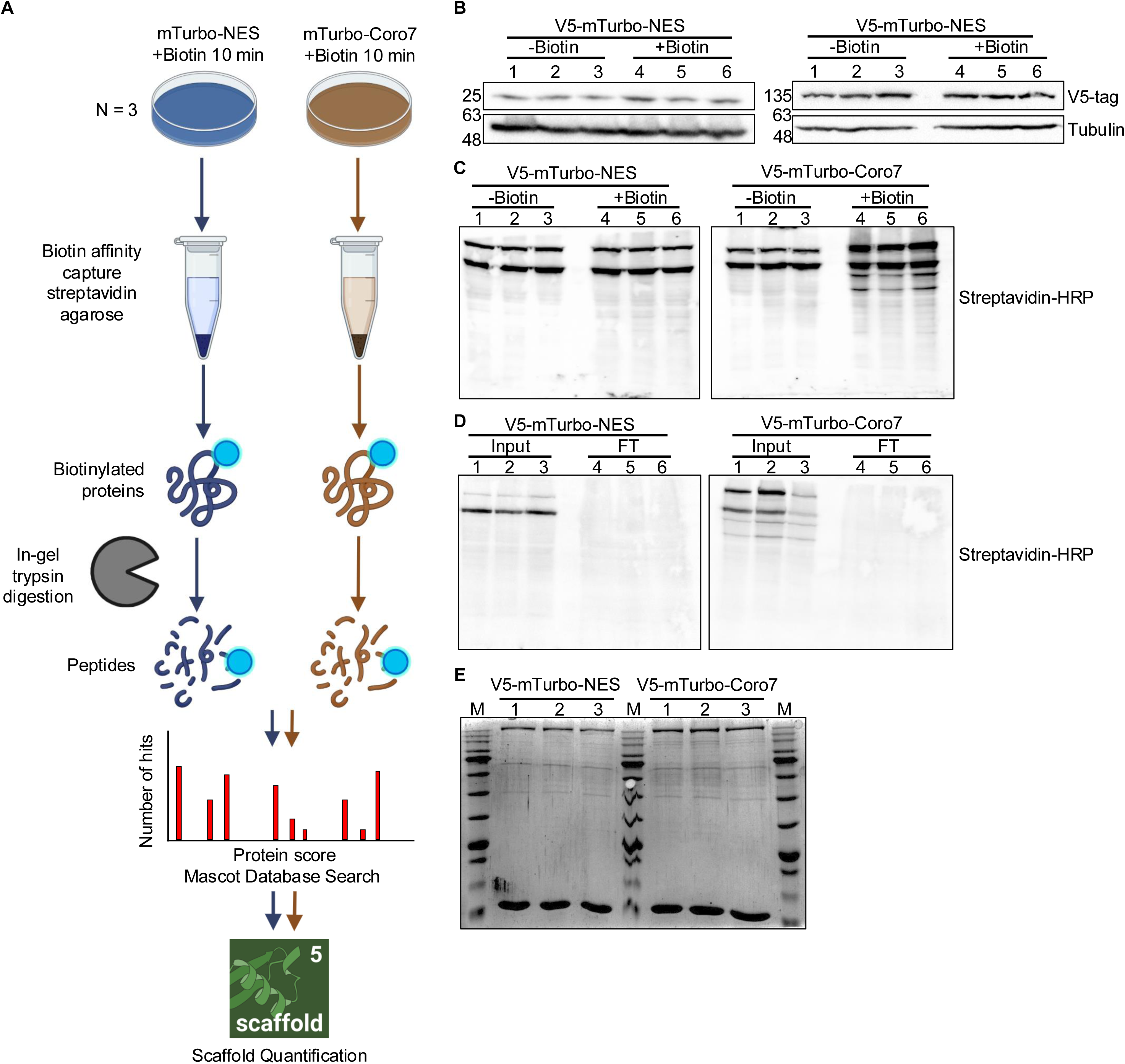
Workflow of V5-mTurbo-Coro7 biotin proximity ligation. **A)** Schematic overview of biotin proximity ligation and mass spec analysis. **B)** Western blot showing equal expression of V5-mTurbo-NES control and V5-mTurbo-Coro7. **C)** Western blot showing biotinylated proteins before and after addition of biotin for V5-mTurbo-NES control and V5-mTurbo-Coro7. **D)** Western blot showing depletion of biotinylated proteins. **E)** Biotinylated proteins shown in C) were concentrated in the stacking gel of a 4-20% stain-free gel, cut out, washed and prepared for mass spec analysis.

